# Evolution at two time frames: polymorphisms from an ancient singular divergence event fuel contemporary parallel evolution

**DOI:** 10.1101/255554

**Authors:** Steven M. Van Belleghem, Carl Vangestel, Katrien De Wolf, Zoë De Corte, Markus Möst, Pasi Rastas, Luc De Meester, Frederik Hendrickx

## Abstract

When species occur in repeated ecologically distinct habitats across their range, adaptation may proceed surprisingly fast and result in parallel evolution. There is increasing evidence that such cases of rapid parallel evolution are fueled by standing genetic variation, but the origin of this genetic variation remains poorly understood. In *Pogonus chalceus* beetles, short- and long-winged ecotypes have diverged in response to contrasting hydrological regimes and can be repeatedly found along the Atlantic European coast. By analyzing genomic variation across the beetles’ distribution, we reveal that genomically widespread short-wing selected alleles evolved during a singular divergence event, estimated at ~0.19 Mya. The ancient and differentially selected alleles are currently polymorphic in all populations across the range, allowing for the fast evolution of one ecotype from a small number of random individuals, as low as 5 to 15, of the populations of the other ecotype. Our results suggest that cases of fast parallel ecological divergence might be the result of evolution at two different time frames: divergence in the past, followed by repeated selection on the divergently evolved alleles after admixture. We suggest that this mechanism may be common and potentially further driven by periods of geographic isolation imposed by large-scale environmental changes such as glacial cycles.

## Introduction

Adaptation to local environmental conditions may lead to the evolution of distinct ecotypes and, ultimately, new species (1, 2). Under prolonged periods of geographical isolation, the absence of gene flow allows populations to accumulate new alleles and build-up genome-wide differences in the frequency of these alleles (3, 4). However, increasing evidence demonstrates that ecological divergence may occur surprisingly fast and even in absence of a physical barrier (5–8). As new beneficial mutations are unlikely to accumulate rapidly, these cases of fast adaptation likely involve selection on genetic variation that was present before divergence took place (9–11). Understanding the origin of and factors that maintain this variation is important as it largely determines the rate and direction of genetic adaptation to rapid environmental change (10, 12, 13).

Populations that have recently and repeatedly adapted to ecologically distinct conditions hold the promise to identify the alleles involved in ecological divergence and to reconstruct their evolutionary history (14–16). Under one scenario, these alleles could have evolved as rare neutral or mildly deleterious alleles within the ancestral population and become repeatedly selected (17). Alternatively, divergently selected genetic variants could initially have evolved and built-up in isolation. Secondary contact between the diverged populations may have then resulted in polymorphisms at these adaptive loci, providing the raw genetic material for repeated and rapid evolution when populations later face similar environmental conditions (18–20). Hence, rapid and repeated ecological divergence could be the result of evolution at two different time frames. This latter idea is appealing as adaptive alleles have in this case already been subjected to a selective filter (10), which may also explain why examples of rapid adaptive divergence often involve parallel adaptation to the same selection pressure. However, complete evidence for this mechanism is currently lacking and requires that (i) differentially selected alleles at multiple unlinked loci diverged within the same time frame, which precedes the more recent parallel divergence events, (ii) differentially selected alleles evolved in at least partial isolation and (iii) populations of each ecotype are polymorphic at these loci to allow for the fast evolution of the alternative ecotype when the selection pressure changes.

Populations of the wing-polymorphic beetle *Pogonus chalceus* have adapted to two contrasting habitat types across Atlantic-Europe; tidal and seasonal salt marshes (Figure 1a). *Tidal* salt-marshes are inundated on an almost daily basis for at most a few hours and are inhabited by *P. chalceus* individuals that have a relatively small body size, short wings and submergence behavior during inundation. In contrast, salt-marshes that are subject to *seasonal* inundations that last for several months harbor *P. chalceus* individuals with a larger body size, fully developed wings and more frequent dispersal behavior (21–23). Populations of both the short- and long-winged ecotypes can be found along the Atlantic coastal region in Europe and often occur in close proximity and even sympatric mosaics (Figure 1b) (24). Despite evidence that ecological divergence is, at least partly, under genetic control (22, 24, 25), previous research based on microsatellite data also revealed very low neutral genetic differentiation within geographic locations (24). This suggests either a very recent differentiation and/or high levels of ongoing gene flow between these ecotypes. These observations render this a most suitable system to infer the origin of the allelic variants that underlie local adaptation (26).

**Figure 1.**
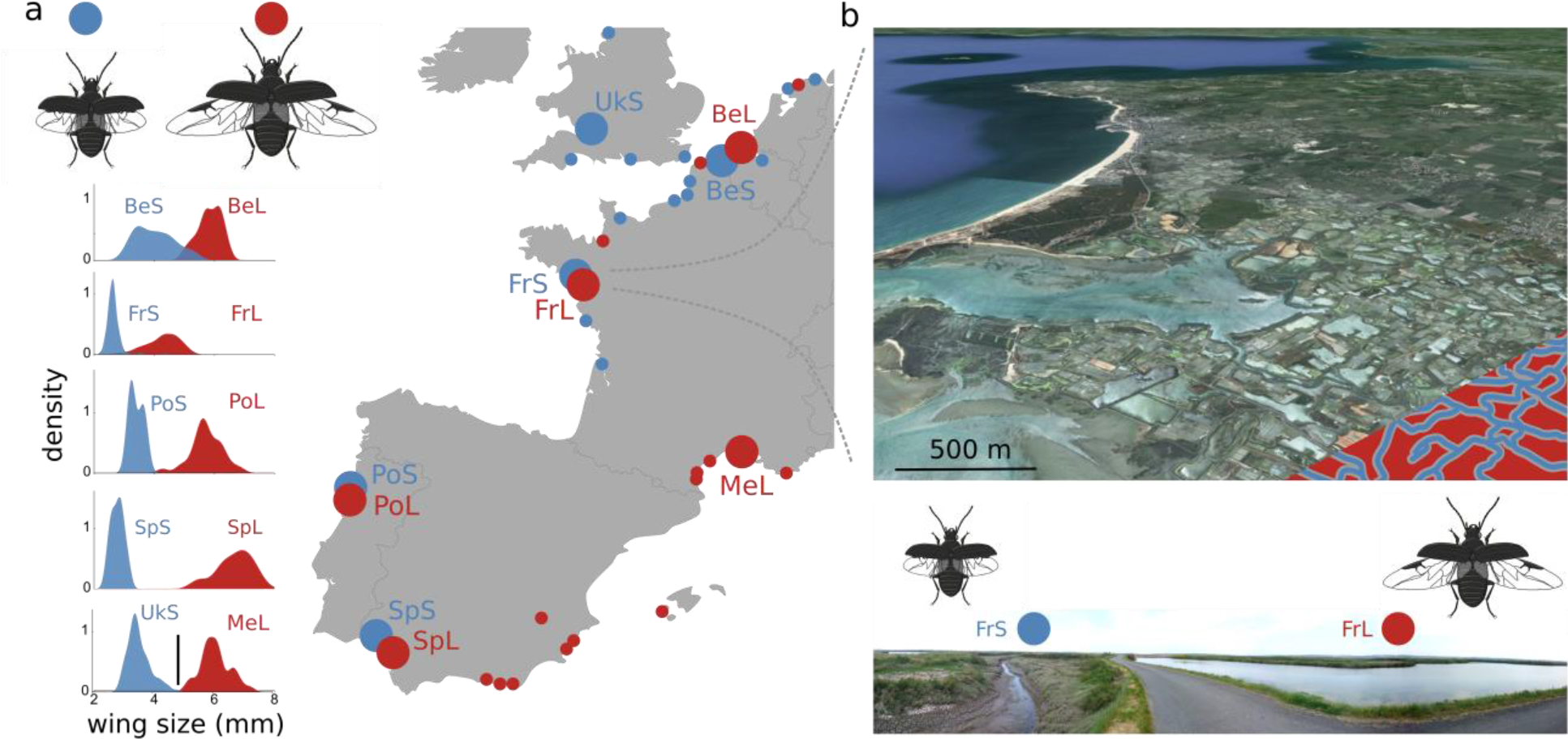
*Pogonus chalceus* sampling and ecotypic divergence. (a.) Sampling locations and density plots of the wing size distribution in the sampled populations. Blue indicates tidal habitats with short-winged beetles, red indicates seasonal habitats with long-winged beetles. BeS: Belgium-short, BeL: Belgium-long, FrS: France-short, FrL: France-long, UkS: UK-short, MeL: Mediterranean-long, PoS: Portugal-short, PoL: Portugal-long, Sps: Spain-short, SpL: Spain-long. Large circles represent populations included in the present study, small circles represent populations sampled previously (22, 31). (b.) Detail of the Fr population (Guérande) where short- and long-winged ecotypes occur in close proximity (< 20 m). This area consists of historic salt-extraction areas that were created by man approximately one thousand years ago and consists of a network of tidally inundated canals (FrS, partly indicated in blue; short-winged populations) interlaced with seasonally inundated salt extraction ponds (FrL, partly indicated in red; long-winged populations).

Here we investigate genomic differentiation in multiple ecologically divergent population pairs of *P. chalceus* and reconstruct the evolutionary history of the alleles underlying ecological divergence. In agreement with a past, singular divergence event, we find sharing of the genealogical pattern at unlinked loci showing signatures of selection. However, the apparent potential of these beetles to rapidly and repeatedly adapt to the different tidal and seasonal hydrological regimes, is likely fueled by the maintenance of relatively high frequencies of the selected alleles in the alternative habitat. These results contribute to our understanding of the mechanisms underlying fast and parallel ecological adaptation and the factors determining the evolutionary potential of populations and species facing changing environments.

## Results

### Wing size distribution

We sampled individuals in four population pairs inhabiting geographically close tidal and seasonally inundated habitats in Belgium (Be; 48 ind.), France (Fr; 48 ind.), Portugal (Po; 16 ind.) and Spain (Sp; 16 ind.), as well as a tidal marsh population in the UK (Uk; 8 ind.) and a seasonally inundated habitat at the Mediterranean coast of France (Me; 8 ind.) (Figure 1a). Individuals from the seasonally inundated habitats had significantly longer wings and larger body sizes compared to those from the tidally inundated marshes (*F*_1,117_ = 1904.4, *P* < 0.0001 for wing length and *F*_1,117_ = 162.29, *P* < 0.0001 for body size). The degree of divergence in wing length between the two ecotypes varied among the four population pairs (*F*_3,117_ = 23.11, *P* < 0.0001), with highly divergent wing lengths in Sp and Po, and some overlap in wing lengths in Be and Fr. Based on these clear-cut differences in wing length, we refer to the populations sampled in the tidal or seasonally inundated habitats as belonging to the short-winged (S) or long-winged (L) ecotype, respectively.

### Population structure and genome wide divergence and diversity

Genetic differentiation (*F_ST_*) among the 10 different populations, based on a quality filtered set of 15,756 SNPs distributed over 1,142 loci obtained by RAD-tag sequencing, varied considerably and ranged from a low (BeS vs. UkS: *F_ST_* = 0.052) to a high degree of differentiation (PoS vs. MeL: *F_ST_* = 0.37) (Table S1). Genetic differentiation increased significantly with increasing geographic distance between the populations (*r_S_* = 0.37, *P* = 0.017) and was higher when populations belonged to a different ecotype (*r_S_* = 0.33, *P* = 0.02) (Figure 2a). When restricting the SNPs to a ‘neutral’ set wherein we excluded RAD-tags containing a SNP with a signature of divergent selection (see *Outlier loci*), there was a stronger effect of geographic distance on genetic differentiation (*r_S_* = 0.54, *P* = 0.002), while the significant ecotype effect disappeared (*r_S_* = 0.09, *P* = 0.2). Principal Coordinate Analysis (PCoA) using all SNP data divided samples largely according to ecotype along the first PCo axis, whereas the second PCo axis grouped samples according to geographic location (Figure 2b). Bayesian clustering (27) of individuals based on their genotypes supported 8 and 6 genetically distinct populations (*K*) for the ‘total’ and ‘neutral’ SNP set, respectively (Figure 2c; Figure S1). For the neutral SNP set, individuals from the same population pair (except Sp) clustered together as a single population, irrespective of their ecotype.

**Figure 2.**
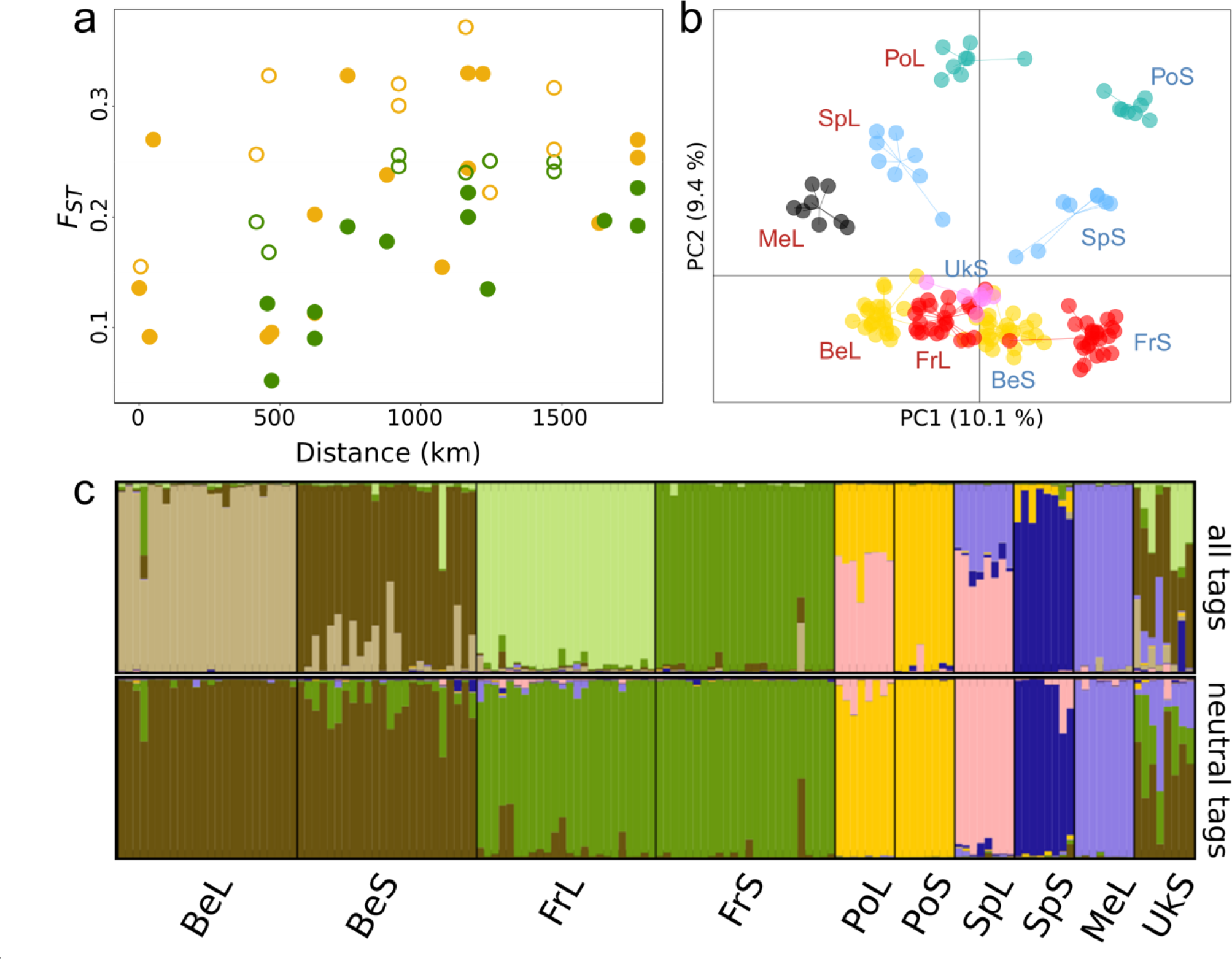
Genetic differentiation among the studied *Pogonus chalceus* populations. **(a.)** Relationship between *F_ST_* and geographic distance. Yellow and green dots depict *F_ST_* comparisons of populations from different or the same ecotype, respectively. Open dots represent *F_ST_* comparisons involving population Po. **(b.)** Principal Coordinate Analysis (PCoA) for RAD-tag sequenced samples. **(c.)** Population structure of the ten *P. chalceus* populations based on Bayesian clustering (27). The best supported number of clusters was 8 for all RAD-tags and 6 for neutral RAD-tags (see Figure S1). See Figure 1 for population codes.

Average nucleotide diversity (*π*) at RAD-tags did not differ between ecotypes (GLMM with RAD-tag ID as random effect: Ecotype effect: *F*_1, 1977_ = 0.31, *P* = 0.6), but differed among population pairs (Population effect: *F*_3, 1977_ = 11.5, *P* < 0.0001). The most southern populations had a significantly higher *π* compared to the northern populations, with Sp having the highest *π* compared to all other populations (P < 0.001) and Po having a higher *π* compared to Fr (*P* = 0.012). The difference in nucleotide diversity among population pairs was also consistent among ecotypes (Population*Ecotype interaction: *F*_3, 1977_ = 2.2, *P* = 0.09).

Despite the apparent close genetic relationship of long- and short-winged ecotypes within each geographic population pair, we observed substantial heterogeneity in *F_ST_* across SNPs (Figure 3, Figure S2). A substantial number of SNPs showed *F_ST_* values that exceeded 0.5 in the ecotype comparisons. For some of these SNPs, different alleles even reached almost complete fixation in the different ecotypes. This proportion of SNPs with *F_ST_* values higher than 0.5 increased towards the more southern population pairs (Be: 2.1%, Fr: 4.5%, Po: 6.3% and Sp: 11.1%). In contrast, only very few *F_ST_* values exceeded 0.5 when similar ecotypes were compared from different population pairs (e.g. Be versus Fr; Figure S2). SNPs that were strongly differentiated in one particular population pair were also significantly more differentiated in any of the other population pairs (0.498 < *r* < 0.66; *P* all < 0.0001; Figure S2), providing support for extensive sharing of highly differentiated SNPs among population pairs.

**Figure 3.**
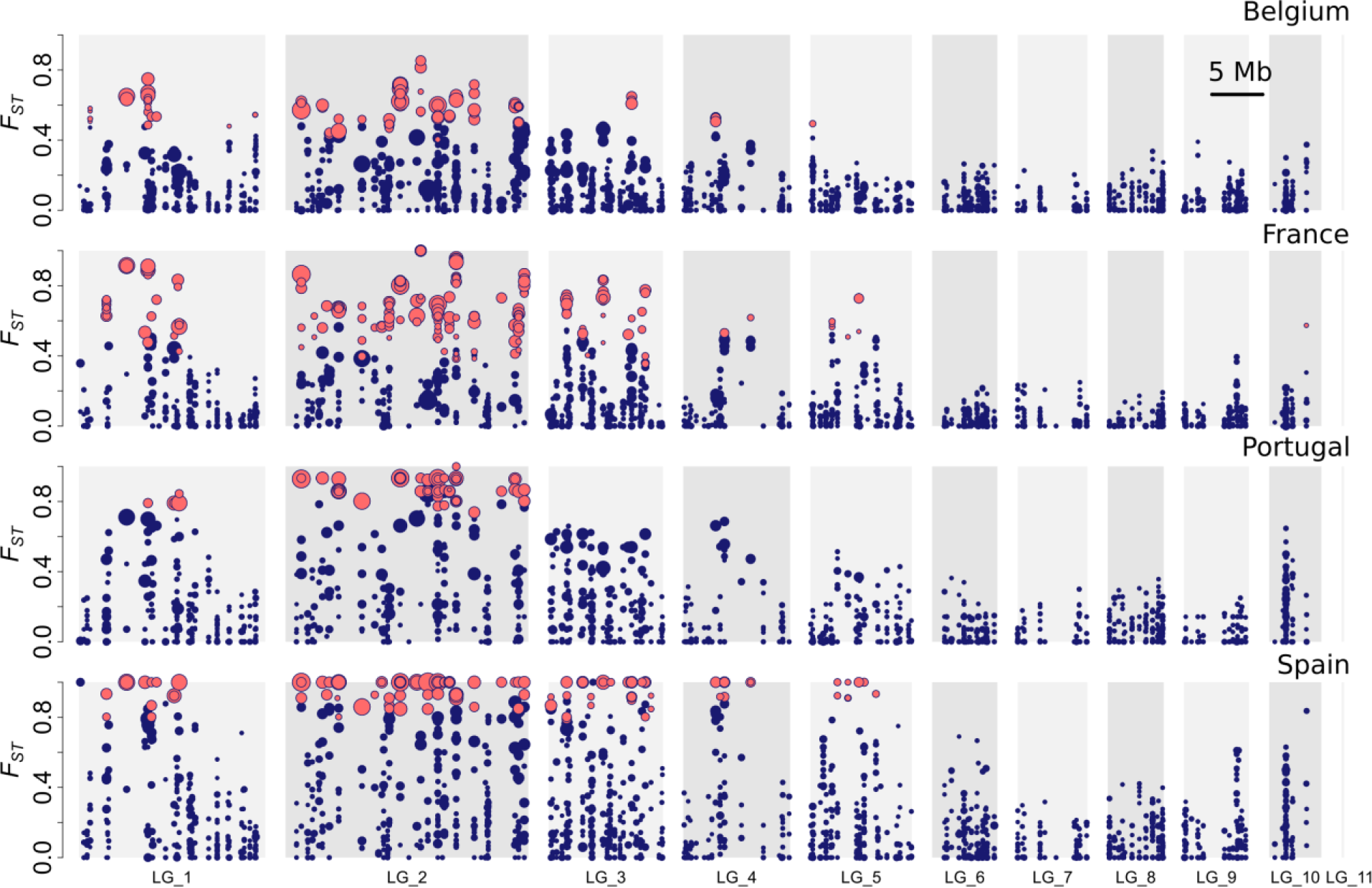
Genomic divergence (*F_ST_*) between *Pogonus chalceus* ecotype pairs. Outlier SNPs within each population pair, as identified by BayeScan (28), are indicated in red. Size of points is proportional to the log_10_BF reported by BayEnv2 (29) and indicates the degree of support that allele frequencies are significantly correlated with habitat type across all sampled populations. Markers on LG10 are significantly sex-linked.

### Outlier loci

Patterns of differentiation were calculated on a per SNP basis and significant outliers were detected using Bayescan (28) and Bayenv2 (29), which allow to account for geographical structure. We identified a total of 512 (3.2%) SNPs that were clustered on 109 (15%) assembled RAD-tag loci with stronger differentiation as expected by chance in at least one of the ecotype comparisons (BayeScan (28) with false discovery rate = 0.05; i.e. on average 4.70 outlier SNPs per RAD-tag locus). A total of 75 (0.48%) SNPs in 32 (6.3%) assembled RAD-tag loci showed allele frequencies that were strongly associated with the ecotypic divergence across all investigated populations (BayEnv2 (29); log_10_BF = 4) (see Supplementary material - outlier loci; Figure S3 and S4). SNPs were mapped to a genome assembly (see Supplementary material – genome assembly and linkage map), showing that SNPs with a high *F_ST_* value clustered into several unlinked regions that were distributed over a large proportion of the genome (Figure 3). These regions with outlier SNPs were largely consistent across the different population pairs and are primarily clustered on the first half of LG1, across the full length of LG2 and LG3 and at the center of LG4 (Figure 3). In contrast, no outlier SNPs were observed on LG6 to LG10. Yet, some more subtle differences could be observed as exemplified by the central region of LG5 where high genomic differentiation was only observed for the Fr and Sp population pair, but not in the Be and Po population comparison. The nuclear-encoded mitochondrial NADP+-dependent isocitrate dehydrogenase *(mtIdh)* locus, that was previously identified to be strongly associated with the ecotypic divergence between short- and long-winged *P. chalceus* beetles (21, 22, 24, 31), is located approximately in the middle of LG2 (scaffold Pchal00589: 148,569-150,947, chromosome LG2: 9,802,630-9,805,108).

### Quantifying standing genetic variation

The diverged blocks on several of the linkage groups likely contain the selected loci of interest (Figure 3). We estimated the potential of each ecotype to adapt to the alternative habitat by calculating the frequency of alternative alleles at outlier SNPs for both ecotypes. Because the identified outlier loci are unlikely to be the exact target of selection, we focused on loci containing SNPs whose allele frequencies were strongly associated with the ecotypic divergence across all investigated populations (BayEnv2 (29); log_10_BF = 4). For each RAD-tag containing multiple SNPs with a BF > 4, only the most strongly supported SNP was selected. Given that these loci are strongly associated with ecotype across all geographic locations, allele frequencies at these loci are expected to be in strong linkage disequilibrium with the allele frequencies at the target of selection. Individuals of the long-winged ecotype contained on average at 10% (SpL) to 42% (BeL) of the outlier SNPs at least one allele associated with the alternative, short-winged ecotype. A similar pattern was found in the opposite direction, with individuals from short-winged populations containing long-wing associated alleles at 10% (SpS) to 48% (BeS) of the outlier SNPs. Moreover, random sampling of genomic variation in an increasing number of individuals showed a steep increase in the proportion of outlier SNPs with at least one allele associated with the alternative habitat (Figure 4). This demonstrates that different individuals from the same population generally carry different alleles that are selected in the alternative habitat at different SNPs. For example, more than 80% of the full set of 75 outlier SNPs detected with BayEnv2 contain at least one short-wing selected allele in a random sample of only eight long-winged individuals for the populations BeL, FrL and PoL (Figure 4a). This demonstrates the presence of substantial standing genetic variation in individuals sampled in the seasonally inundated marshes to adapt to tidal marshes. Only for the most southern long-winged populations (SpL and MeL), outlier loci that are likely linked to short-wing selected alleles are present at lower frequencies and these long-winged populations are unlikely to contain the full set of alleles selected in the short-winged ecotype. For FrS, PoS and particularly SpS, long-wing selected alleles accumulated at a much lower rate under random sampling of short-winged individuals (Figure 4b).

**Figure 4.**
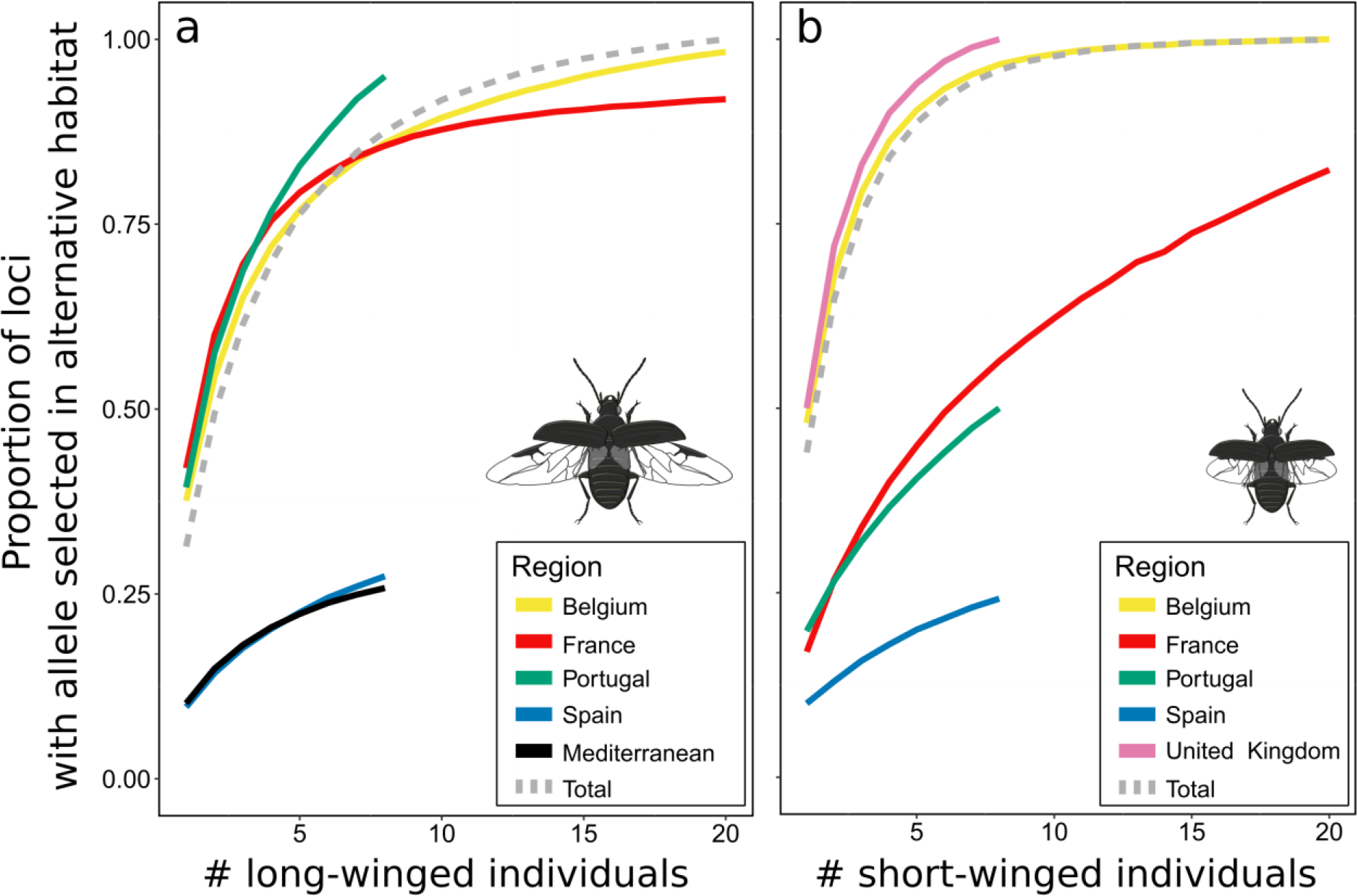
Amount of standing genetic variation. Accumulation curves of **(a.)** the proportion of outlier loci containing at least one short-wing selected allele in a random sample of *N* long-winged individuals and **(b.)** the proportion of outlier loci containing at least one long-wing selected allele in a random sample of *N* short-winged individuals.

### Sequence variation and phylogenetic reconstruction at outlier loci

Haplotype networks and trees of the 1.2 kb sequence alignments obtained from RAD-tag loci show that haplotypes selected in short-winged populations generally clustered as a strongly supported monophyletic clade of closely related sequences (clade support level > 0.96; Figure 5a, b and S5). This clustering supports a singular mutational origin of short-wing selected alleles at each of the investigated outlier tags. Short-wing selected alleles appeared to be derived as they most frequently constituted a subclade within those selected in long-winged populations (Figure S5). This is in line with the observation that all other species within the genus *Pogonus* are long-winged (32). The average absolute divergence between the differentially selected haplotypes (*d_XY_* = 0.011 ± 0.0014) was about 1.65 times higher compared to the average divergence between two randomly chosen haplotypes at these loci (*π*_*tot*, outliers_ = 0.0067 ± 0.00097, *t*-test: *P* < 0.0001) and highlights a deep divergence between the alleles selected in short- and long-winged populations. Dating the divergence time between the differentially selected alleles using BEAST (33) and the divergence from *P. littoralis* as a calibration point (0.62 Mya (31)), pointed towards highly comparable divergence times among outlier loci (Figure 5b and S5). The divergence time between long- and short-wing selected alleles ranged between 0.12 Mya and 0.28 Mya, with an average of 0.189 Mya ± 0. 09 Mya, and suggests that the divergence of short-winged alleles from the long-winged alleles took place during the Late Pleistocene.

**Figure 5.**
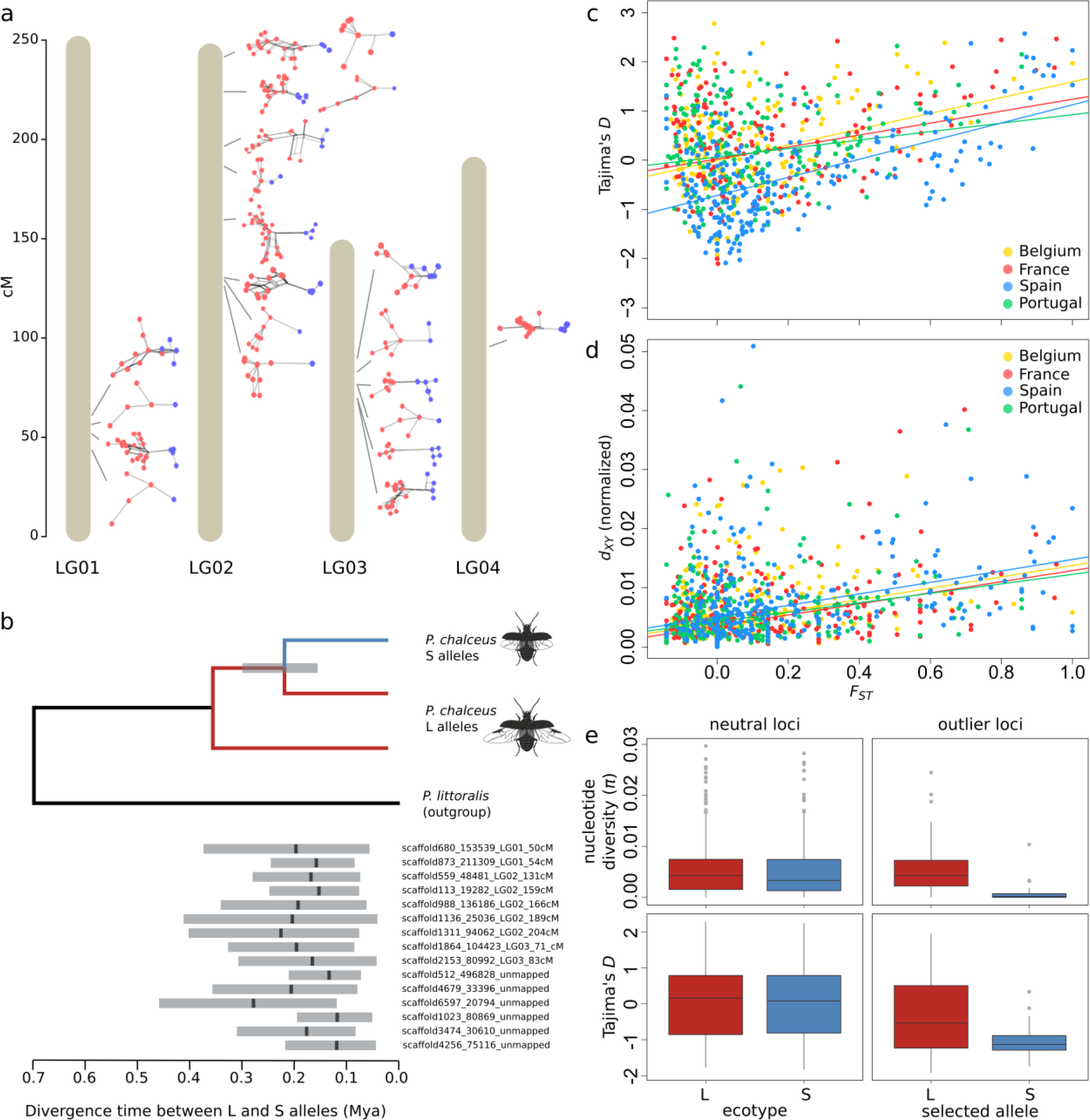
Haplotype structure and diversity at divergently selected loci. (a.) Haplotype networks of RAD-tags containing outlier SNPs and at least 10 variable sites (BayEnv2; log_10_BF > 4) at the different linkage groups. Haplotypes selected in short-winged populations are depicted in blue, haplotypes selected in long-winged populations are depicted in red. (b.) Estimated divergence time (Mya) between alleles selected in short-winged (blue) versus long-winged (red) populations. The tree represents the general phylogenetic relationship between short- and long-wing selected alleles and the estimated divergence point. (c.) Relationship between *F_ST_* and Tajima’s *D* and (d.) absolute nucleotide divergence, *d_XY_*, scaled relative to the divergence from the outgroup species *Pogonus littoralis* in the four population pairs. (e.) Comparison of nucleotide diversity (*π*) and Tajima’s *D* at neutral loci of long-winged (L) and short-winged (S) populations (left) and between haplotypes at outlier RAD-tags selected in L or S populations (right).

In agreement with divergent selection, sequences from more strongly differentiated RAD-tags had a significantly higher Tajima’s *D* (i.e. increased heterozygosity; *F_ST_*: *F* = 57.36, *P* < 0.0001; Figure 5c) and absolute nucleotide divergence between the ecotypes normalized by the divergence from the outgroup *P. littoralis* (= *d_XY_ / d_XY, P. littoralis_*, see Methods for details; *F_ST_*: *F* = 83.9, *P* < 0.0001; Figure 5d). The significant relation between *F_ST_* and absolute divergence (normalized *d_XY_*) further supports that the observed heterogeneity in genomic divergence between the ecotypes is the result of divergent selection of ancient alleles embedded within a genome that is homogenized between the ecotypes, rather than selection on recently obtained new mutations (34, 35). Furthermore, a reduced recombination rate was observed between haplotypes that are divergently selected between long- and short-winged populations (*r*^2^ = 0.140) compared to the recombination rate observed within long-winged haplotypes (*r*^2^ = 0.184).

We further observed that nucleotide diversity (*π*) of haplotypes selected in the short-winged ecotype was strongly reduced and tended to be nearly seven times lower (*π*_S_ = 0.0009 ± 0.0003) compared to those selected in the long-winged ecotype (*π*_L_ = 0.0062 ± 0.0003) (GLMM with tagID as random effect: Ecotype effect: *F*_1_, 66 = 25.12, *P* < 0.0001; Figure 5e). This difference was consistent among the four populations (Ecotype*Population interaction: *F*_3_, 64 = 0.87; *P* = 0.45). In contrast, nucleotide diversity at neutral RAD-tags was comparable between both ecotypes (GLMM with RAD-tag as random effect: Ecotype effect: *F*_1_, 599 = 0.98, *P* < 0.3; Figure 5e). The nucleotide diversity of the haplotypes selected in long-winged populations was also comparable to average nucleotide diversity observed at non-outlier loci (*π*_tot, neutral RAD-tags_ 0.0057 ± 0.0003), showing that only haplotypes selected in the short-winged ecotype have this reduced nucleotide diversity (Figure 5e). Similarly, Tajima’s *D* of haplotypes selected in the short-winged ecotype was significantly lower compared to those selected in the long-winged ecotype (*F_1,27_* = 11.7; *P* = 0.002; Figure 5e) and suggests a recent spread of short-wing selected alleles along the Atlantic European coast.

## Discussion

Understanding the genomic basis of repeated and fast ecological adaptation provides unique insights into the process of evolutionary diversification (14, 15, 36). While evidence is accumulating that cases of repeated adaptation are largely driven by selection on standing genetic variation (37), the evolutionary origin of the selected alleles remains currently poorly understood (20, 38). In *P. chalceus*, several unique observations help to disentangle the complex history of fast and parallel ecological divergence and the origin of the adaptive alleles that underlie it. These observations include: (i) Diverged ecotypes within a geographic region share genomic variation at neutral loci but are characterized by extensive genomic islands of divergence, widely distributed across the genome. (ii) Genomic regions of elevated divergence are highly consistent across the four population pairs. (iii) Haplotypes of alleles selected in short-winged populations cluster monophyletically and are derived. (iv) The pattern of divergence across short-wing selected alleles and divergence time estimates are comparable among loci. (v) Short-wing associated haplotypes show strongly reduced diversity, negative Tajima’s *D* and reduced recombination with the long-wing associated haplotypes. (vi) A relatively small subsample of individuals from long-winged populations harbor the majority of alleles associated with short-winged populations, and vice versa. In summary, these observations imply that the ecotypes diverged *in situ* within each geographic region, but that the adaptive alleles that constitute the short-winged ecotype have a singular evolutionary origin and likely evolved within an ancient subpopulation. Moreover, the monophyletic clustering of short-wing selected haplotypes and their deep divergence and reduced recombination with long-wing selected haplotypes suggest that this short-winged subpopulation evolved in isolation from the long-winged populations. The time at which short-wing selected alleles diverged was estimated to be about 0.19 Mya. Subsequent admixture between the long- and short-winged subpopulation may then have resulted in a genetically polymorphic population from which the short-winged ecotype re-evolved more recently and repeatedly along the European Atlantic coast by the concerted selection of these previously diverged alleles.

Despite the ancient divergence of the adaptive alleles, the contemporary repeated distribution of the *P. chalceus* ecotypes likely evolved from highly admixed populations. The low amount of neutral genetic differentiation should, however, not be interpreted as conclusive evidence for a repeated *in situ* divergence, because it also may be the result of neutral gene flow between both ecotypes at secondary contact (38–40). Nonetheless, several additional observations support that the current distribution of the ecotypes involves the repeated *in situ* evolution of both ecotypes. First, quantifying the amount of short-wing selected alleles present in long-winged populations revealed that sufficient genetic variation is present in a founding population of between 5 and 15 long-winged individuals to evolve a short-winged ecotype. Thus, genetic constraints for the evolution of the short-winged ecotype out of long-winged individuals, and vice versa, appear to be surprisingly low. Both weak selection as well as high rates of ongoing gene flow between the ecotypes could explain the relatively high frequency of alleles that are selected in the alternative ecotype (41). Second, short-winged individuals are unable to disperse by flight between the currently highly isolated salt-marsh areas. The fragmented distribution of tidal salt marshes along the Atlantic coast renders it therefore unlikely that they were colonized by short-winged individuals based on terrestrial dispersal alone, in particular because the species strongly avoids unsuitable habitat patches (32). Third, we previously put forward a behavioral mechanism that may explain the spatial sorting of these ecotypes into their respective habitats (i.e. long-winged beetles tend to avoid frequent flooding in tidal habitats, whereas short-winged beetles stay submerged during short tidal flooding events), which may reduce gene flow and induce *in situ* divergence of the genetically distinct ecotypes (23). Adding to this, our data agree with an initial divergence of multiple genetically unlinked loci in geographic isolation, which may have further facilitated the repeated ecological divergence under gene flow (42).

Major geographic expansions and contractions of the tidal and seasonal habitat types and the *P. chalceus* species distribution have likely occurred since the initial divergence of the short-winged ecotype. We estimated the evolution of the short-winged ecotype to have occurred during the Mid to Late Pleistocene. Since then, Europe has been subject to at least one interglacial (0.130 – 0.115 Mya) and one glacial (0.115 – 0.012 Mya) period. These major climatic changes fragmented the Euro-Atlantic coastline, potentially creating opportunities for the initial evolution of the short-winged ecotype in the partially isolated large coastal floodplains that extended, for instance, around the North-Sea basin (43). Suggested by the negative Tajima’s *D* and low nucleotide diversity among the short-wing selected alleles, the short-winged ecotype likely spread along the Atlantic European coasts from a small refuge population and came into secondary contact with the long-winged populations more recently, after the last glacial period. During the last glacial maximum, the refuge population was likely located more southward relative to the current species distribution, as it is deemed unlikely that the species persisted at the current northern latitudes of its distribution (32). In addition, the lower degree of overall genetic differentiation between the ecotypes in the more northern population pairs Be and Fr, together with their higher frequencies of alleles selected in the alternative habitat, less profound phenotypic differentiation and lower overall genetic diversity (*π*) are all consistent with a northward expansion of a highly admixed and dispersive long-winged population and more recent ecotypic divergence. Potentially, this expansion may have further facilitated the maintenance of deleterious short-wing selected alleles in the expanding long-winged population (44), which then became repeatedly selected in the emergent tidal (short-winged) habitats.

The two-step process of initial divergence in an ancient and potentially isolated population and subsequent admixture putatively also applies to other examples of fast and repeated ecological divergence. Repeated ecological divergence at the same loci has been reported in some iconic examples of parallel evolution, such as stickleback, cichlid fishes and *Heliconius* butterflies (45–50). These loci have in many cases been assigned to shared ancient polymorphisms that were present in the population before the evolution of the currently observed divergent populations (50). Moreover, many of these loci are sometimes identified and are unlinked throughout the genome, such as in fruit flies, *Timema* walking sticks and *Littorina* sea snails (7, 51, 52). The genetic signature of the evolution of the *P. chalceus* ecotypes shows strong analogies to these well-studied cases of repeated adaptation. In cichlid fishes, moreover, it has been extensively argued that divergence in isolation and subsequent admixture may have provided the genetic material for the incredibly diverse and recent adaptive radiations of cichlid fish (11, 53). Untangling the evolutionary history of the alleles involved in these and other cases will help in better understanding the processes that drive parallel divergence as well as fast responses to environmental change

## Conclusion

The initial evolution of co-adapted alleles at multiple physically unlinked loci is facilitated in geographic isolation (3). Subsequent admixture of gene pools may then enrich the adaptive genetic variation and allow for subsequent fast and repeated adaptation. In agreement to this, in *P. chalceus* populations the alleles required to adapt to the alternative environment are maintained in the source population. These loci are expected to be maladaptive within the source population and it is likely both the temporal and spatial repetition of this divergence, combined with relatively high levels of gene flow and range expansion, that maintain these allele frequencies. Glacial cycles, in particular, can be expected to have played an important role in this process as episodes of fission and fusion of the different ecotypes generates strong opportunities for both the evolution of adaptive genetic variants as well as admixture (43). Historic selection pressures could therefore play a pivotal role in determining the rate, direction and probability of contemporary adaptation to changing environmental conditions.

The proposed mechanism illustrates that the distinction between *in-situ* divergence and secondary contact is less clear-cut as generally assumed if populations are highly admixed and that both processes can be involved at different time frames. An important implication is that this mechanism might reconcile different views on the geography of ecological divergence in which adaptive divergence between closely related populations is either interpreted as primary divergence, and thus the onset of speciation (2), or the result of secondary introgression after initial ecological divergence in allopatry (38–40).

## Methods

### Sampling

Diverged population pairs of *P. chalceus* were collected from both tidal and seasonal salt marshes extending nearly the entire species range (Figure 1) (32). We sampled four geographically isolated population pairs (separated between approximately 450 km and 900 km) of a tidal and seasonally flooded inland population each. Wing and elytral sizes were measured by means of a calibrated ocular with a stereomicroscope. We further conducted RAD-seq genotyping on two specimens of the long-winged outgroup species *P. littoralis*, which were sampled in the Axion Delta, Thessaloniki, Greece (Table S2).

### RAD-tag sequencing

DNA was extracted using the DNA extraction NucleoSpin^®^ Tissue kit (Macherey-Nagel GmBH). Extracted genomic DNA was normalized to a concentration of 7.14 ng/µl and processed into RAD libraries according to Etter et al. (2011), using the restriction enzyme SbfI-HF (NEB) (54). Final enrichment was based on 16 PCR cycles. A total of nine RAD libraries including 16 individuals each and, hence, a total of 144 individuals were sequenced paired-end for 100 cycles (i.e. 100 bp) in a single lane on an Illumina HiSeq2000 platform according to manufacturer’s instructions. The outgroup *P. littoralis* specimens were sequenced separately. The raw data was demultiplexed to recover individual samples from the Illumina libraries using the *process_radtags* module in Stacks v1.20 software (55). Reads were quality filtered when they contained 15 bp windows of mean Phred scores lower than 10. PCR duplicates were identified as almost (i.e. allowing for sequencing errors) identical reverse read sequences and removed, using a custom Perl script (56).

### Genome assembly

Total DNA was extracted from individuals captured in the canal habitat of the salt marshes in the Guérande region (France), using the DNA extraction NucleoSpin® Tissue kit (Macherey-Nagel GmBH). Illumina paired-end (100 bp) and mate-paired (49 bp) libraries were constructed with insert sizes of 200 bp, 500 bp, 800 bp, 2 kb and 5 kb and sequenced on an Illumina HiSeq2000 system according to the manufacturer’s protocol (Illumina Inc.). Adapter contamination in reads was deleted using Cutadapt v1.4 (57) and reads that did not have a matching pair after adaptor filtering were removed. Reads were corrected for sequencing error with SOAPec v2.02 (58), using a *k*-mer size of 17 and a low frequency cutoff of consecutive k-mer of 3. Sequencing of the 200 bp, 500bp, 800 bp, 2 kb and 5 kb insert libraries resulted in a total of ~57.7 Gb of sequencing data, of which 56.6 Gb was retained after data cleaning (Table S3). Reads were assembled using SOAPdenovo2 (58) using a *k*-mer parameter of 47, which was selected for producing the largest contig and scaffold N50 size after testing a range of *k*-mer settings between 19 and 71. The short insert libraries were used for both contig building and scaffolding. The long insert libraries were only used for scaffolding. SOAPdenovo GapCloser v1.12 tool (58) was used with default settings to close gaps emerging during scaffolding. We used DeconSeq v0.4.3 (59) to identify and remove possible human, bacterial and viral contamination in the assembly (Table S4). Completeness of the assembled genome was assessed by comparing the assembly with a dataset of highly conserved core genes that occur in a wide range of eukaryotes using the CEGMA pipeline v2.5 (60).

### Linkage map

To position the genomic scaffolds into linkage groups, we constructed a linkage map by genotyping parents and offspring (RAD-seq) from four families (Table S5). For the parental generation, we used lab-bred individuals (*F*_0_) whose parents originated from the French population, to ensure that they had not been mated in the field. A total of 72 *F*_1_ offspring (n = 23, 14, 23 and 12 offspring from each family) were raised till adulthood and subsequently genotyped, together with their parents. To maximize the number of scaffolds comprising a marker, RAD-tag sequencing was based on a PstI-HF (NEB) digest (6 bp recognition site) instead of the SbfI-HF (NEB) digest (8bp recognition site) of the population genomic analysis. Final enrichment was based on 16 PCR cycles. Illumina HiSeq sequencing resulted in a total of 237 M paired-end reads, of which 113 M remained after quality filtering and removal of PCR duplicates. Reads were mapped to the draft reference genome with BWA-mem (61) using default settings. Linkage map reconstruction was performed with LepMAP2 (62). LepMAP2 reconstructs linkage maps based on a large number of markers and accounts for lack of recombination in males due to achiasmatic meiosis, which is suggested in *P. chalceus* and male Caraboidea in general (63) (see Supplementary material – genome assembly and linkage map).

### Population genomic analysis

Quality and clone filtered paired-end reads of the 144 field captured individuals were mapped to a draft reference genome with BWA-mem (61). Indel realignment, SNP and indel calling was performed with GATK’s UnifiedGenotyper tool (64). Paired-end sequencing of the approximately 200 to 600 bp RAD tag fragments adjacent to symmetric SbfI restriction sites allowed us to obtain sequence information of 1,200 bp fragments (paired RADtag) around each restriction site. Hence, after SNP calling we retained all sites within 1,200 bp windows around each SbfI recognition site in the genome, totaling 732,884 bp of sequence. Haplotype phasing was subsequently first performed with GATK ‘read-backed phasing’ (64), while the remaining unphased sequences were phased with Beagle v4.1 (65). A reliable SNP set was then obtained by filtering out SNPs with minimum genotype quality lower than 20, minimum average depth lower than 10 and a minor allele frequency less than 0.01. This resulted in 27,757 SNPs, distributed over 712 paired RADtags, with an average individual depth of 62.9 (± 51 std), of which 15,756 were present in at least 80% of the individuals.

### Analysis of population structure

Pairwise *F*_ST_-statistics (66) across RAD-tags were calculated for all pairwise population comparisons using Genepop v4.5.1 (67). Principal Coordinate Analysis was performed using *adegenet* in R (68). To minimize dependence due to physical linkage among SNPs, we randomly selected one single SNP per paired RAD-tag. We also constructed a ‘neutral’ subset by excluding SNPs located on scaffolds showing signatures of divergent selection (20.5% of all SNPs). As a criterion, we excluded scaffolds containing a SNP with a log_10_(BF) >3 as determined by BayEnv (69). The Pearson correlation between genetic divergence and either geographic distance between the populations or ecotype (coded as 1 = different ecotype and 0 = identical ecotype) was assessed by a Mantel test in the vegan v2.2-1 package in R v3.1.3 (70). Based on these two datasets, we used the Bayesian clustering algorithm implemented in STRUCTURE v2.3.4. (27) to assign individuals into *K* clusters based on their multilocus genotype. We applied an admixture model with three independent runs for each *K* = 2-10, 100,000 MCMC repetitions with a burn-in of 30,000, correlated allele frequencies among populations and no prior information on population origin. Default settings were used for the prior parameters. The best supported number of clusters (*K*) was determined from the increase in the natural logarithm of the likelihood of the data for different numbers of assumed populations.

### Outlier loci detection

Support for loci showing significantly higher degrees of differentiation was first detected with BayeScan2.1 (28) within each population pair (Be, Fr, Po and Sp). BayeScan assumes that divergence at each locus between populations is the result of population specific divergence from an ancestral population as well as a locus specific effect. The prior odds of the neutral model was set to 10. Twenty pilot runs, 5,000 iterations each, were set to optimize proposal distributions and final runs were performed for 50,000 iterations, outputting every tenth iteration, and a burn-in of 50,000 iterations. Detection of outlier loci is particularly vulnerable to false positives (71). To account for this, we applied a false discovery rate (FDR) correction of 0.05, meaning that the expected proportion of false positives is 5% (28).

To test for the presence of SNPs whose alleles are directionally selected in the two habitats across all populations, we used the approach implemented in BayEnv2 (29, 69). This method identifies SNPs whose allele frequencies are strongly correlated with an environmental variable given the overall covariance in allele frequencies among populations. The covariance in allele frequencies, which represents the null model against which the effect, *β*, of an environmental variable on the allele frequencies of each SNP is tested, was estimated based on all SNPs present in at least 80% of the individuals. This covariance matrix was strongly correlated with the *F_ST_* matrix (Mantel-test: *r_S_*= 0.87), indicating that it accurately reflects the genetic structuring of the populations. For each SNP, the posterior probability of a null model assuming no effect of the environment (**β** = 0) is compared against the alternative model which includes the effect of the environmental variable. As environmental variable, we assigned tidal habitats (BeS, FrS, PoS, SpS and UkS) the value −1 and seasonal inundated habitats (BeL, FrL, PoL, SpL and MeL) the value 1. The degree of support that variation at a SNP covaries with the habitat wherein the population was sampled is then given by the Bayes Factor (BF), the ratio of the posterior probabilities of the alternative versus the null model. For both the estimation of the covariance structure and the environmental effect, a total of 100,000 iterations was specified.

### Reconstructing the evolutionary history of outlier loci

To gain insight into the evolutionary history of the alleles differentiating the two ecotypes, we reconstructed haplotypes of the 1200 bp long paired RAD-tag loci for each individual. Sites with a read depth lower than 10 or a genotype quality lower than 20 were treated as missing. Haplotypes could be reconstructed for 627 paired RAD-tags with on average 671 bp genotyped in at least 75 % of the individuals. We constructed split networks with the NeighbourNet algorithm using SplitsTree4 (72) for all RAD-tags that contained an outlier SNP with a Bayes Factor (BF) support level larger than 3 based on the BayEnv2 analysis. Haplotypes were subsequently split in two groups according to base composition at the outlier SNP with the highest support and visualized on the networks.

We further calculated for all RAD-tags the following haplotype statistics with the EggLib v2.1.10 Python library (73): haplotype based *F_ST_* (30), average pairwise difference (*d_XY_*) between both ecotypes, total nucleotide diversity (*π_tot_*), nucleotide diversity within the long- and short-winged ecotype (*π_L_* and *π_S_*, respectively) and Tajima’s D. Comparison of measures of *d_XY_* (≈ 2*μ*t + θ_Anc_) and *π* (≈ 4*Nμ*) between RAD-tags depend, besides the average coalescence time between haplotypes, also on the mutation rate (*μ*) of the RAD-tag. As we are primarily interested in comparing values of these statistics among RAD-tags independent of their mutation rate, we normalized these values by the average number of nucleotide differences between *P. chalceus* and the outgroup species *P. littoralis* (74). More specifically, we first calculated *d_XY_* between haplotypes of *P. chalceus* and *P. littoralis* (*d*_*xy*, littoralis_), and divided both *π_tot_* and *d_XY_* by this value.

We estimated the divergence time between haplotypes selected in short- and long-winged populations with BEAST 1.7.1 (33). The analysis was restricted to outlier RAD-tags (15 in total) that are also present in the outgroup species *P. littoralis* and that contained on average at least 10 segregating sites among the sequences of *P. chalceus*. This latter criterium was implemented to ensure a sufficiently high substitution rate for reliable time calibration. The tree was calibrated using the divergence from *P. littoralis*, estimated at 0.62 ± 0.8MY, as calibration point (31). We assumed a GTR substitution model, a strict clock model and standard coalescent tree prior. Analyses were run by default for 10 million generations of which the first 2 million generations were treated as burn-in and discarded for the calculation of posterior probability estimates.

## Data accessibility

Raw sequencing reads are available in the NCBI Short Read Archive under BioProject PRJNA381601. The genome assembly, ordered using the linkage map, is available under accession NEEE00000000. Reads of genome assembly: SAMN06684244-SAMN06684249; RAD-seq data of *Pogonus chalceus*: SAMN06691389-SAMN06691532; RAD-seq data of *Pogonus littoralis:* SAMN06691533-SAMN06691534; RAD-seq data for linkage map construction: SAMN06806679-SAMN06806758.

## Acknowledgments

This study could not have been performed without access to *P. chalceus* collection of the late Konjev Desender. We are grateful to Alexandre Ramos, Viki Vandomme and Lut Van Nieuwenhuyse for help in collecting the samples, Jonas Van Belleghem for measuring wings and Simon Martin for valuable bioinformatics help, Karim Gharbi (Edinburgh Genomics Center) for extensive support in the preparation of the RAD-seq libraries and Chris Jiggins for valuable comments on the manuscript. This work was supported by funding received from the FWO-Flanders (PhD grant to SVB) and the Belgian Science Policy (MO/36/ 025, BR/121/PI/GENESORT and BR/175/PI/PARAWINGS to FH) and was partly conducted within the framework of the Interuniversity Attraction Poles program IAP (SPEEDY) – Belgian Science Policy. The genomic analyses were carried out using the STEVIN Supercomputer Infrastructure at Ghent University, funded by Ghent University, the Flemish Supercomputer Center (VSC), the Hercules Foundation and the Flemish Government – Department EWI.

## Author contributions

SMVB and FH designed the research and analyzed the data. SMVB, CV, KDW and FH performed the research. SMVB, FH, CV, ZDC, MM and LDM wrote the paper. PR contributed analytical tools.

